# The Hidden Side of Diversity: Effects of Imperfect Detection on Multiple Dimensions of Biodiversity

**DOI:** 10.1101/2021.06.02.446400

**Authors:** Aline Richter, Gabriel Nakamura, Cristiano Agra Iserhard, Leandro da Silva Duarte

## Abstract

1. Studies on ecological communities often address patterns of species distribution and abundance, but few consider uncertainty in counts of both species and individuals when computing diversity measures.
2. We evaluated the extent to which imperfect detection may influence patterns of taxonomic, functional, and phylogenetic diversity in ecological communities.
3. We estimated the true abundance of fruit-feeding butterflies sampled in canopy and understory strata in a subtropical forest. We compared the diversity values calculated by observed and estimated abundance data through the hidden diversity framework. This framework evaluates the deviation of observed diversity when compared with diversities derived from estimated true abundances, and if such deviation represents a bias or a noise in the observed diversity pattern.
4. The hidden diversity values differed between strata for all diversity measures, except for functional richness. The taxonomic measure was the only one where we observed an inversion of the most diverse stratum when imperfect detection was included. Regarding phylogenetic and functional measures, the strata showed distinct responses to imperfect detection, despite the tendency to overestimate observed diversity. While the understory showed noise for the phylogenetic measure, since the observed pattern was maintained, the canopy had biased diversity for the functional metric. This bias occurred since no significant differences were found between strata for observed diversity, but rather for estimated diversity, with the canopy being more clustered.
5. We demonstrate that ignore imperfect detection may lead to unrealistic estimates of diversity and hence, to erroneous interpretations of patterns and processes that structure biological communities. For fruit-feeding butterflies, according to their phylogenetic position or functional traits, the undetected individuals triggered different responses in the relationship of the diversity measures to the environmental factor. This highlights the importance to evaluate and include the uncertainty in species detectability before calculating biodiversity measures to describe communities.

## Introduction

Estimating the whole biodiversity in a community is a key challenge for ecologists. First, because we do not have time and resources to sample all species and individuals that are present in a community. Second, even focusing on a target group, there are large proportions of species or individuals that remain “hidden” (Chao et al., 2017; Devarajan, Morelli, & Tenan, 2020; Guillera-Arroita, Kéry, & Lahoz-Monfort, 2019; Yoccoz, Nichols, & Boulinier, 2001). This occurs since both species and individual are not perfectly observed in the field (i.e. they are undetected during sampling), and different species have distinct probabilities of being detected (Boulinier, Nichols, Sauer, Hines, & Pollock, 1998; Ribeiro, Williams, Specht, & Freitas, 2016). Classical community analyses commonly ignore imperfect detection, for both incidence and abundance-based approaches, as well as its effects on diversity measures (DeVries, Alexander, Chacon, & Fordyce, 2012; Pillar & Duarte, 2010; Wiens & Donoghue, 2004). Identify the effects of imperfect detection in classical diversity measures might improve our understanding of relationships between diversity and environmental gradients (Roth, Allan, Pearman, & Amrhein, 2018), and ultimately the processes that structure the biological communities (Dorazio, Connor, & Askins, 2015).

A considerable portion of community studies that employed models that account for imperfect detection (e.g., Multi-Species Hierarchical Models) are interested in evaluating the true occurrence or abundance, aiming to guide management practices (Ruiz-Gutiérrez, Zipkin, & Dhondt, 2010; Yamaura et al., 2012; Zipkin, Andrew Royle, Dawson, & Bates, 2010). But, the effects of imperfect detection are not restricted only to the taxonomic aspect of diversity (e.g. species richness), and our ability in detecting biodiversity patterns may differ among different components of diversity (Iknayan, Tingley, Furnas, & Beissinger, 2014; Jarzyna & Jetz, 2016). Species co-occurring in communities exhibit different levels of shared evolutionary history and variation in phenotypic traits. These features of species are widely used to infer historical and/or ecological mechanisms determining community assembly patterns (Duarte, Debastiani, Carlucci, & Diniz-Filho, 2018; Graham & Fine, 2008; Webb, Ackerly, McPeek, & Donoghue, 2002). Despite the increase in studies that quantified phylogenetic or functional diversity (de Bello et al., 2015; Tucker et al., 2017), few consider the imperfect detection in species count for calculate it (Chao et al., 2017; Frishkoff, De Valpine, & M’Gonigle, 2017) or have quantified the role and magnitude of the effects of imperfect detection on distinct facets of diversity (Jarzyna & Jetz, 2016; Si et al., 2018). If undetected species have unique phylogenetic information or functional traits, by underestimating their contribution to diversity estimate, we are neglecting an ecologically important part of the assemblages (Jarzyna & Jetz, 2016). Consequently, we would observe a more clustered assemblage than they really are (Si et al., 2018). The opposite can also occur when undetected species are phylogenetically or functionally redundant (Jarzyna & Jetz, 2016), and the observed assemblages will overestimate phylogenetic and functional diversity. Furthermore, the detection of species can be biased at some part of the environmental gradient evaluated (Roth et al., 2018). If this occurs, not only the observed diversity pattern can be affected, but also our interpretation of the relationship among diversity and environmental gradients (Fig. 1a-b).

**Figure 1.**
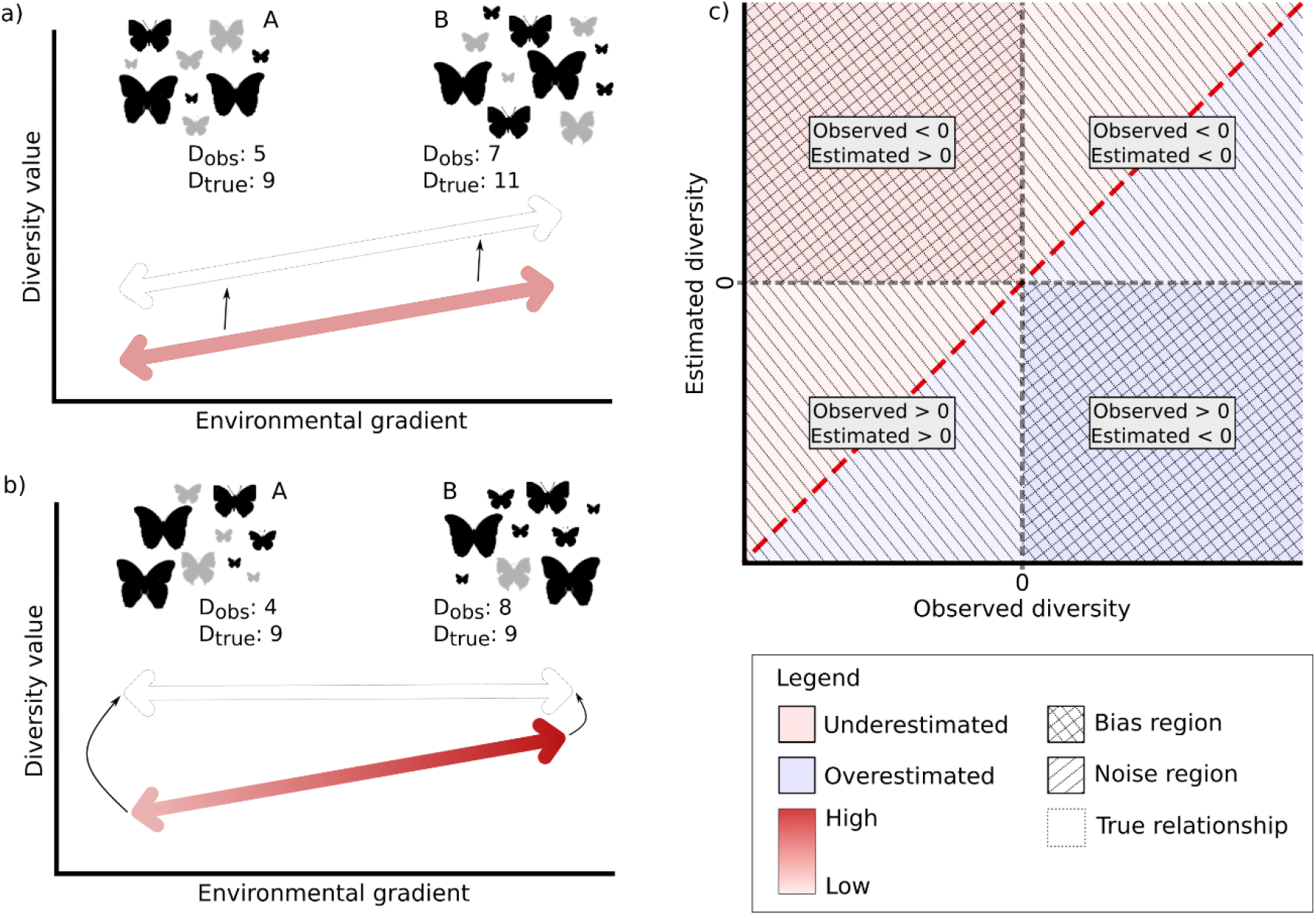
Schematic representations of the hidden diversity framework. a) and b) represents the relationship between diversity values and environmental gradients. c) demonstrate in which way the imperfect detection can influence this relationship. In a) each set of butterflies represents a community, called A and B. D_obs_ is a value of a diversity measure calculated from data of an observed community containing imperfectly detected species (detection probability (p) < 1), and D_true_ represents the real value of that given diversity if all species in the community were sampled (p = 1). The difference between the true and the observed values represents the hidden diversity (HD). For community A the hidden diversity is the same along the environmental gradient, HD = −4. Despite the lack of 4 unities of a given measure (low detectability, filled red arrow), community B remained more diverse than A, i.e., the pattern was maintained despite the noise yielded by imperfect detection (true relationship, dashed hollow arrow). In b) we also have two communities, A and B. Here, there is a difference between HD values for A and B (−5 and −1, respectively), and the relationship among diversity measure and the environmental gradient is biased by imperfect detection: observed diversity in B is higher than A because diversity is more detected in B than A (red gradient arrow), but in fact, they have the same value of diversity (D_true_ = 9). However, despite the differences in a) and b) scenarios, both have the observed diversity underestimated in relation to the truth, represented by red quadrants in the upper diagonal for c). This visual evaluation of hidden diversity allows us to observe if underestimated values were yielded by an inversion in the observed pattern (e.g., from clustered to overestimated, or vice-versa). When this occurs, the points will fall in the top-left or bottom-right right checkered quadrants, which we defined as critical bias regions, because even if there no is difference along the environmental gradient, therefore the pattern remains the same, the inference of processes that assembly the community will be wrong. Hachured quadrants indicate noise regions because even if the observed diversity is largely affected by imperfect detection, the observed pattern will not change, but still can vary according to an environmental factor. The dashed red line represents the hidden diversity, points above it indicate underestimated diversity and have negative HD values, while points under it indicate an overestimation of diversity and have positive HD values.

Insects are the most species-rich taxa in the world, which poses a major challenge for ecologists interested in evaluating insect diversity patterns (Thomas, 2005). Among insect groups, butterflies are considered important biological indicators due to their short life-cycle and high sensibility to changes in environmental features (New, 1997, Brown & Freitas, 2000). Fruit-feeding butterflies are a conspicuous guild of tropical butterflies that feed on rotting fruit, carrion, or plant exudates (DeVries, 1988) and represent about 50 – 75% of nymphalid diversity in the Neotropical region (Brown, 2005). Assemblages of fruit-feeding butterflies show high vertical stratification (Devries, 1988; DeVries, Alexander, Chacon, & Fordyce, 2012; Ribeiro & Freitas, 2012; Santos, Iserhard, Carreira, & Freitas, 2017), with the canopy generally being taxonomically more diverse than understory. These strata exhibit a large difference in their microclimatic conditions, habitat structure and, hence in their taxonomic composition (Araujo, Freitas, Souza, & Ribeiro, 2020; DeVries et al., 2012; Santos et al., 2017). Whereas Charaxinae, Biblidinae, and Nymphalinae are recognized as canopy-dwellers, Satyrinae is generally associated with understory sites (Schulze, Linsenmair & Fiedler, 2001). In a phylogenetic or functional perspective, the composition of those strata could be also dissimilar, once that lineages of fruit-feeding butterflies exhibit habitat preferences (Fordyce & DeVries, 2016) and individuals tend to show traits that varying according to characteristics and preferences (Graça, Pequeno, Franklin, & Morais, 2017).

Due to their feeding habit, these butterflies can be sampled with passive and standardized methodologies such as bait traps (Freitas et al., 2021). Unlike other methods to sample butterflies (entomological nets or transect counts), bait traps avoid bias related to variation in the observer or personal expertise about species detection (Boulinier et al., 1998, Kéry & Plattner, 2007, Ribeiro et al., 2016). However, the detection of individuals might be biased by bait attractiveness in different habitats and by the individual ability to find the trap. Weather conditions as wind speed, rain, and temperature, can influence the bait volatiles, leading to decreased attractiveness, especially in open habitats (Marini-Filho & Martins, 2010). Fruit-feeding butterflies typically use odor cues to locate food, and some groups, such as Charaxinae, are able to find more accurately their preferred food (Molleman, Alphen, Brake, & Zwaan, 2005). Further, individuals that have high mobility, may often be undetected in a sampling site because it is visiting other sites within their home range (Joseph, Elkin, Martin, & Possingham, 2009). Therefore, bearing in mind the intrinsic challenges of sampling in the canopy together with the characteristics of individuals that inhabit this stratum, it is more likely that the canopy has a higher number of undetected individuals than understory, yielding a bias in diversity measured in this stratum.

In this study, we aimed to analyze the extent to which imperfect detection, assessed by the estimates of the true abundance of species, can lead to changes in observed patterns of taxonomic, functional, and phylogenetic diversities of butterflies living in different forest strata (canopy *vs.* understory). We expect that: (i) canopy will show lower individual detection than understory, leading to a source of bias that hides the true diversity value for this stratum. Consequently, this bias induces an erroneous inference when we compare diversity values between canopy and understory. (ii) The effect of imperfect detection will be lower for phylogenetic and functional measures concerning taxonomic diversity. In this case, an increment in species number will not be followed by an increment in both phylogenetic and functional diversity, indicating that undetected species are redundant with species sampled in the observed community.

## Material and Methods

### Study sites and sampling procedures

The study site was located in Floresta Nacional de São Francisco de Paula (FLONA-SFP; centered at 29°25’22’’S, 50°23’11’’W) in Southern Brazil. FLONA-SFP comprises an area of 1,615 ha in the Atlantic Forest biome and is composed of Mixed Ombrophilous Forest with the presence of *Araucaria angustifolia* (Bertol.) Kuntze, as well as patches with *Pinus* sp. and *Eucalyptus* sp. plantations (ICMBio, 2020). The climate of the region is temperate without a dry season, and with annual mean rainfall near 2,000 mm and an annual average temperature of 14.5°C (Sonego, Backes, & Souza, 2007).

Fruit-feeding butterfly assemblages were sampled between November 2016 and March 2017, which correspond to the summer season in the Southern Hemisphere and which is the best period of the year for sampling butterflies in the Atlantic Forest (Iserhard, Romanowski, Richter, & Mendonça, 2017). We adopted standardized methods for sampling fruit-feeding butterflies in the Neotropical region (Freitas et al., 2014), which consisted in install five traps per sampling unit, which were baited with a mixture of mashed banana and sugarcane juice (Freitas et al., 2021). We performed monthly surveys at six sites of native forest within FLONA-SFP for five months. In each month, the traps remained open for eight to ten consecutive days and every 48h the traps were checked and the bait was replaced. This totalizes a sampling effort of 2,520 trap-days (10 traps × 6 sampling units × 42 sampling days). In each site, we sampled the assemblages of fruit-feeding butterflies in the canopy (~15 m above the ground, inside canopy tree crowns) and in the understory (1.5 m above the ground) and each stratum was considered as one independent sampling unit. In every trap checking, we measured the temperature of the base of each trap using an infrared thermometer (GM-300, Benetech®).

### Community model for abundance data

We employed a modification of the Dorazio-Royle-Yamaura model (DRY) (Kéry & Royle, 2016; Yamaura et al., 2011; Yamaura, Kéry, & Andrew Royle, 2016) to estimate uncertainties in the individual counts for fruit-feeding butterflies. The modifications allow the model to estimate the mean abundance (λ_ik_) and detection probability (p_ijk_) for each stratum (Zipkin et al., 2010). We assumed that local abundance remained unchanged during the survey (i.e. closure assumption, Kéry, Royle, & Schmid, 2005) since we sampled in a narrow time window, and that mean abundance and detection probability were independent among species. Abundance for each species *k* at each site *i* is a latent variable (i.e. imperfectly observed) called *N_ik_*, which follows a Poisson distribution:

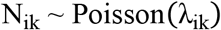

where *λ_ik_* is the mean or expected abundance. We assumed that *λ_ik_* varied among sites depending on species random effects and if point *i* was in the canopy (Strata = 0) or the understory (Strata = 1), thus allowing species-level effects to differ between the two strata (Zipkin et al., 2010). We also included a slope for the mean temperature obtained from the base of the traps of each site *i* (Temp) and add two random site effects, because samplings were repeated in time (sampling months, SM) for each sampling units (SU), and hence their measures are not independent within them. We fit the model for biological process using a log-link function, as follows:

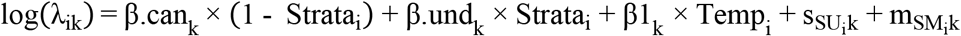

where β.can and β.und are the species-specific intercepts for canopy and understory, respectively, β1 is the species-specific slope for the temperature effect, *s* and *m* are the random effects for six sampling units and five sampling months.

We describe the detection process as:

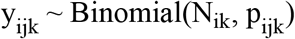

where the number of detected individuals *y_ijk_* during visit *j* was obtained with *N_ik_* trials and a probability of successful detection *p_ijk_*. The detection history *y_ijk_* > 0 indicates that the species *k* (1, 2, …, 35) was observed in site *i* (1, 2, …, 12) during the sampling occasion *j* (1, 2, …, 5), while *y_ijk_* = 0 implies the species was undetected. We modeled detectability as a logit-linear combination of species-specific detection probabilities dependent on the stratum and two covariates:

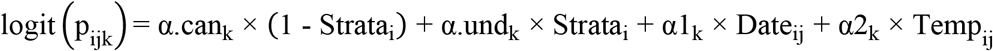

where α.can and α.und are the species-specific intercepts for canopy and understory, respectively, and α1 is the linear effects of the sampling day (transformed to Julian date) and α2 is the linear effects of the temperature by day.

All covariates for the biological and observation process were standardized before perform the Bayesian model. The effect of predictors was corroborated whenever 95% of the Credible Interval (CRI) did overlap zero. We defined species-specific parameters for each stratum and for covariates as coming from normal hyper-distributions, e.g., β.can_k_ ~ Normal (μβ._can_, τβ._can_), being that these priors describe the heterogeneity among species. We determined vague priors for the hyper-parameters that define the mean (μ) and precision (τ) at the community-level, such that μ ~ Normal (0, 0.001) and τ, that is the inverse of variance (τ = sd^−2^), where sd ~ Uniform (0, 10), and these hyper-parameters are shared by all species in each stratum (Yamaura et al., 2016). Considering that the mean detection probability must vary between 0 and 1, we defined μα = logit(μ_α.pre_), when μ_α.pre_ ~ Uniform (0, 1), and then, α_k_ ~ Normal (μ_α_, τ_α_). The model was run using the package *jagsUI* (v. 1.4.9, Kellner, 2015) with three Markov Chains Monte Carlo (MCMC), 150,000 iterations with the first 50,000 iterations discarded, and a thinning rate of 100. The model code is given in Appendix A (N-mixture model). These settings of MCMC results in a posterior sampling with 3,000 iterations. We also defined initial values for parameter N and monitored the community mean and species-level parameters. We checked the convergence of MCMC by R-hat statistics (Gelman & Rubin, 1992) and graphical visualization.

In addition, we checked and validated the N-mixture model through simulation of metacommunities. For each simulation, we set the mean expected abundance for canopy and understory (β_s1_ and β_s2_) or the mean probability for canopy and understory (α_s1_ and α_s2_) to vary, while all other parameters were kept constant. For each parameter, we defined true mean values, which we consider low, intermediate, and high, resulting in 12 simulated metacommunities (hereafter treated as setting code A to L). The output of the simulation provided two main information: the true abundance of species for each community (N_s_) and the imperfect observed community (yobs_s_). The yobs_s_ was then subjected to the N-mixture model, and we monitored all parameters estimated. For the biological model, all true values of parameters and hyper-parameters fall within 95% of the credible interval of the posterior distribution (Appendix B – Fig. B1 to B3), indicating that model was able to recovery true parameters values.

### Phylogenetic and functional data

We collected at least one specimen of each butterfly species captured in bait traps for subsequent measurement of functional traits. We selected 12 functional traits to characterize functional diversity in each community, including traits related to flight performance, habitat use, and ecological behavior (Table 1) (Chai & Srygley, 1990; Dudley, 2002; Spaniol, Duarte, Mendonça, & Iserhard, 2019). Using the recently proposed phylogeny of Chazot et al. (2019) for Nymphalidae, we obtained the phylogenetic relationships among the 35 species of fruit-feeding butterflies recorded in this study. We pruned the complete tree to calculate measures of phylogenetic diversity and structure of communities. We used the packages *ape* (v. 5.3, Paradis & Schiliep, 2019), and *phytools* (v. 0.6-44, Revell, 2012) to prune the tree.

**Table 1.** Description for the functional traits measured for fruit-feeding butterflies sampled at FLONA-SFP, southern Brazil. C – continuous traits, B – binary traits.

|  | Trait name | Type | Measure | Description | References |
| --- | --- | --- | --- | --- | --- |
| FWL | Forewing length<br>(mm) | C | Forewing base to apex | Used as a proxy for body size and<br>related with dispersion capacity | Chai and Srygley<br>1990, Sekar 2012 |
| TM:TDM | Thorax mass to<br>total body mass<br>ratio | C | The ratio between thorax<br>mass and total body mass | The proportion that represents the<br>investment in thorax mass; related to<br>flight capacity due that thorax allocates<br>the flight muscles | Chai and Srygley<br>1990 |
| AM:TDM | Abdomen mass to<br>total body mass<br>ratio | C | The ratio between abdomen<br>mass and total body mass | The proportion that represents the<br>investment in abdomen mass; related to<br>investment in reproductive tissues | Srygley and Chai<br>1990 |
| FEA | Functional eye area (mm<sup>2</sup>) | C | Set of linear eye measurements | Represent the functional visual field; associated with habitat perception | Rutowski 2000, Turlure et al. 2016 |
| WL | Wing loading (N/m<sup>2</sup>) | C | Amount of body mass sustained by wing area unit | Related with flight speed and agility and can be associated with adaptative response to environmental gradients | Chai and Srygley 1990, Berwaerts et al. 2002, Turlure et al. 2016 |
| AR | Aspect Ratio | C | The ratio between forewing span squared to forewing area | Express the wing shape; related to flight speed and agility | Chai and Srygley 1990, Berwaerts et al. 2002 |
| FS | Food specialization | C | Amount of host plants used by immature stages | Express the food habit; lower values represent specialists and higher values represent more generalists species. | Graça et al. 2017 |
| Iridescence | Wing Iridescence | B | Presence or absence of iridescence coloration | Related with intra and interspecific visual recognition | Pinheiro et al. 2016, Spaniol et al. 2019 |
| Eyespots | Wing Eyespot | B | Presence or absence of wing eyespots | Related with defense strategies to avoid or deflect attacks of visually hunting predators | Stevens 2005, Olofsson et al. 2010 |
| Rings | Mimetic Ring | B | Member or not of mimetic rings complex | Indicate if species are a member of mimetic rings; related to Mullerian, Batesian or scape mimetic rings | Su et al. 2015, Spaniol et al. 2019 |
| Camouflage | Camouflage strategies | B | Colorations and shapes that resemble background or environmental structures | Related to capacity to avoid predators | Ruxton et al. 2004; Skelhorn et al. 2010 |
| Disruptive | Disruptive Coloration | B | Conspicuous colorations in the wing's periphery that disguises the body outline of the animal | Related to capacity to avoid predators, by preventing prey recognition | Schaefer and Stobbe 2006 |

### Incorporating imperfect detection in diversity measures: The Hidden Diversity framework

To evaluate the magnitude of the effects of imperfect detection on diversity measures we developed an R function called *hidden.diversity* (HD) (Appendix C – Hidden diversity framework). This function returns, for each site *i*, the deviation of observed diversity from the estimated diversity, given imperfect detection, and this difference is divided by the standard deviation of the estimated diversity as follow:

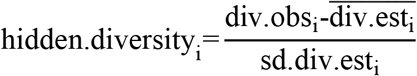

where *div.obs_i_* is the taxonomic, functional or phylogenetic diversity value obtained with observed count data for each site, 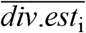 is the mean diversity value obtained from *N_ik_* posterior sampling in each site and *sd.div.est_i_* is the standard deviation of *div.est_i_*. Positive and negative values of HD indicate, respectively, an overestimation and underestimation of observed diversity in relation to estimated diversity values. Overestimation of diversity can only occur for phylogenetic or functional measures, since that species included can be functionally or phylogenetically redundant, and the N-mixture model only accounts for false negatives. However, distinct scenarios can generate positive or negatives HD values, we disentangle these possible scenarios by plotting the relationship between observed and estimated diversity values (Fig. 1c). We called noise when observed and estimated diversity has the same signal, in other words, the observed pattern (overdispersion or clustering) does not change after corrected by imperfect detection, but still can be overestimated or underestimated in comparison with the estimated true diversity. On the other hand, a bias will occur if the observed and estimated diversity have opposite signs, and for these cases, an erroneous pattern in phylogenetic and/or functional structure of communities will be observed when undetected species are not considered.

The input of the *hidden.diversity* function is the observed community data, a phylogenetic tree, a matrix containing the mean traits for each species, and the matrix *N_ik_* estimated by the N-mixture model which represents the detection-corrected abundance. The function internally calculates taxonomic diversity (TD), the standardized effect size for phylogenetic diversity (SES.PD), and functional diversity (SES.FD), as well as the SES for phylogenetic and functional structure (SES.MPD and SES.MFD respectively). Also, the function allows indicating if there are binary data in the trait matrix, if the diversity measures should be weighted by abundance, the type of null model, and the number of permutations used to calculate the null models, which aimed to remove the effect of species richness on diversity measures. Null models were built by randomizing communities by permuting species positions on the phylogenetic tree or functional dendrograms, or by permuting sampling units (rows) or species identities (columns) in the community matrix. Null models are implemented in the package *picante* (Kembel et al., 2010). The function output is a data frame containing SES values of diversity measures for each site (observed and estimated) and the value of hidden diversity.

We employed the HD for each diversity measure to evaluate differences between canopy and understory in the bias yielded by imperfect. For this, we performed a linear mixed model (LMM) using the HD values for each diversity measure as the response variable, the strata as a fixed predictor, and the sampling months and sites as random factors.

## Results

Our database contained 35 species and 914 individuals of fruit-feeding butterflies. We found that canopy had lower community-level mean abundance than understory (values in the natural scale, μ_β.can_ = 0.166 CRI_95%_ = 0.008 to 0.104, μ_β.und_ = 2.655, CRI_95%_ = 0.001 to 0.117). Moreover, understory assemblages had a higher mean detection probability (μ_α.can_ = 0.032, CRI_95%_ = 0.025 to 0.038, μ_α.und_ = 0.497, CRI_95%_ = 0.033 to 0.964) (Fig. 2). We do not explore the effects of predictor variables on abundance and detection probability because these results are not crucial for this study, but the values for hyper-parameters for community-level are shown in Appendix A (Fig. A1 and A2, Table A1).

**Figure 2.**
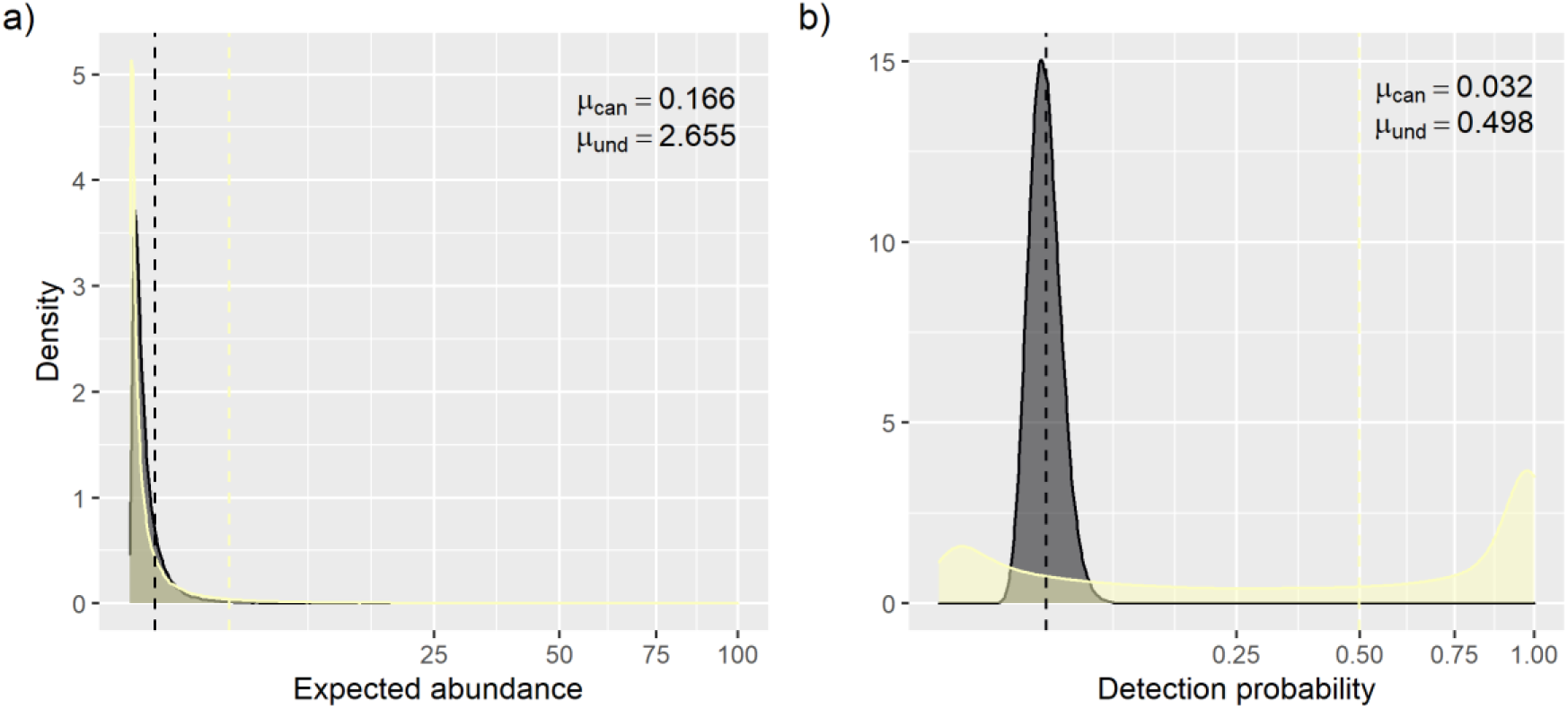
Community-mean distribution for expected abundance (a) and detection probability (b) for fruit-feeding butterflies sampled at FLONA-SFP, Southern Brazil. These distributions were generated using the community hyperparameters for canopy (μ_can_ and sd_can_, black color) and understory (μ_und_ and sd_und_, light-yellow color). The dashed line represents the mean for each stratum (μ). We apply a square root transformation on the *x*-axis to better improve the visualization.

Hidden diversity (HD) demonstrated that there was an underestimation for both strata when only the richness was evaluated (TD), and for this diversity measure, the HD did not differ between strata (Fig. 3a, Table 2). All other diversity measures tended to be overestimated (positive HD values). Phylogenetic and functional measures had opposite responses in relation to the most overestimated stratum, while for phylogenetic measures understory was more overestimated than the canopy, for functional measures canopy tended to show higher overestimation than understory. Only for SES.FD we did not observe a difference in the HD between strata (Table 2). However, observing the relationship among observed and estimated diversity, we found that for most sites, the pattern of positive or negative SES value was maintained. This implies that, despite the error associated with not accounting for imperfect detection, for the fruit-feeding butterfly assembly, imperfect detection acts more like a noise than a bias (Fig. 3b).

**Figure 3.**
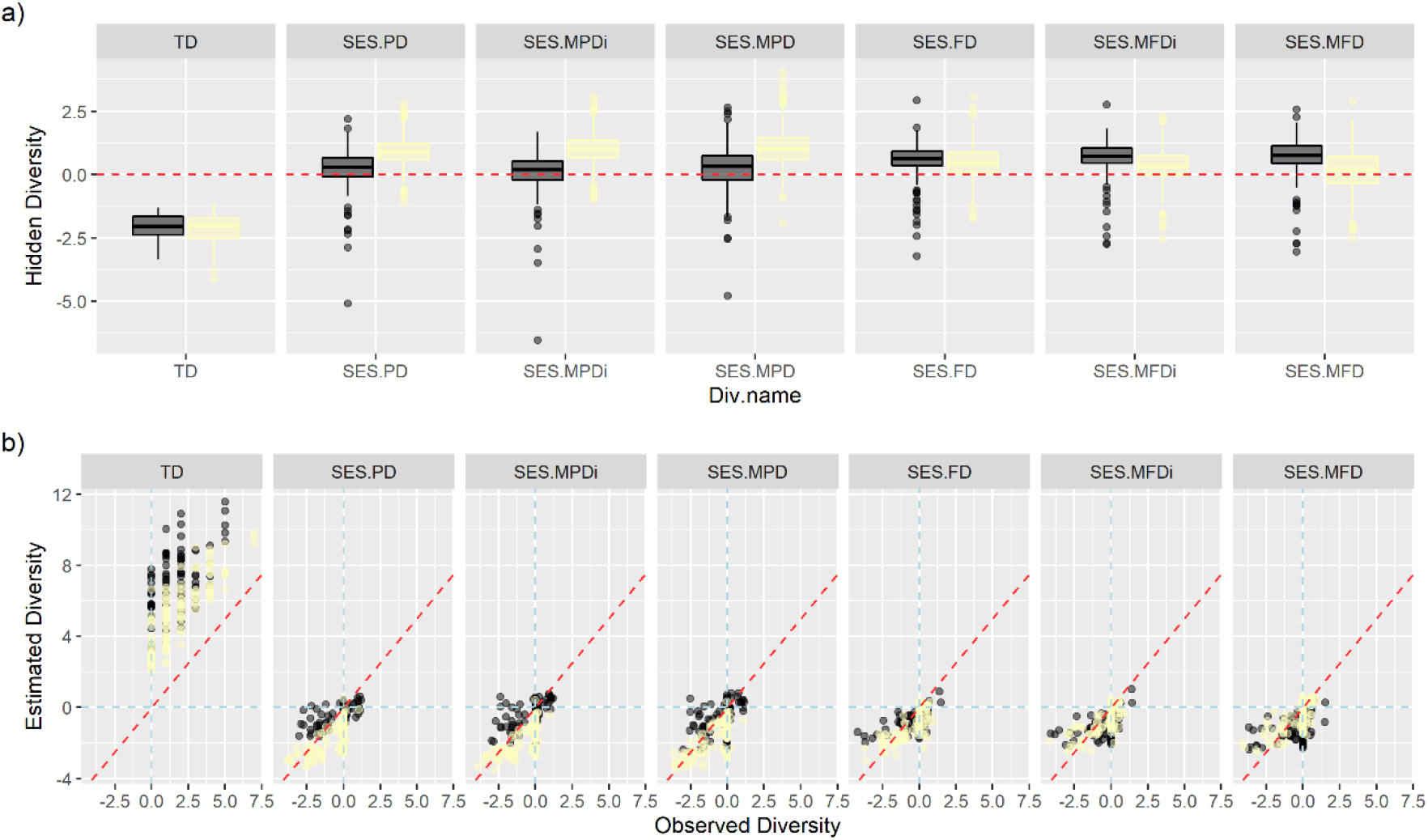
The effects of imperfect detection on multiple dimensions of biodiversity, evaluated by the hidden diversity framework, for an assemblage of fruit-feeding butterflies sampled at FLONA-SFP, southern Brazil. a) Response of each stratum – Canopy (dark boxplots) and Understory (yellow boxplots) – to the imperfect detection and their variation among the diversity measures. TD – taxonomic diversity, SES – standardized effect size, PD/FD – phylogenetic/functional diversity, MPD/MFD – abundance-based mean pairwise phylogenetic/functional distance, MPD_i_/MFD_i_ – incidence-based mean pairwise phylogenetic/functional distance. The red dashed line indicates no difference in diversity value between observed and estimated data. b) Visual evaluation of the effect of the imperfect detection by sampling unit (points) and environmental gradient (colors, dark – Canopy; yellow – Understory). Points above the dashed red line indicate an underestimate of the diversity and negative values of hidden diversity; points below this line indicate an overestimation of diversity and positive values of hidden diversity.

**Table 2.** Relationship of hidden diversity values for taxonomic, phylogenetic, and functional measures (HD.TD, HD.PD/MPD, HD.FD.MFD) and vertical stratification for the assemblage of fruit-feeding butterflies sampled at FLONA-SFP, southern Brazil. Bold values indicate a statistical significance at a threshold of 0.05. The asterisk indicates values of mean pairwise distances calculated with incidence instead of abundance.

|  | Estimate | SE | t value | p |
| --- | --- | --- | --- | --- |
| **H.TD** |  |  |  |  |
| intercept | -2.064 | 0.163 | -12.679 | **0.000** |
| Slope | -0.099 | 0.046 | -2.163 | **0.031** |
| **H.PD** |  |  |  |  |
| intercept | 0.174 | 0.090 | 1.931 | 0.092 |
| Slope | 0.728 | 0.090 | 8.060 | **0.000** |
| **H.FD** |  |  |  |  |
| intercept | 0.509 | 0.064 | 7.955 | **0.000** |
| Slope | -0.032 | 0.090 | -0.349 | 0.727 |
| **H.MPD<sub>i</sub>\*** |  |  |  |  |
| intercept | 0.076 | 0.128 | 0.595 | 0.572 |
| Slope | 0.938 | 0.095 | 9.849 | **0.000** |
| **H.MPD** |  |  |  |  |
| intercept | 0.226 | 0.177 | 1.279 | 0.240 |
| Slope | 0.848 | 0.104 | 8.164 | **0.000** |
| **H.MFD<sub>i</sub>\*** |  |  |  |  |
| intercept | 0.619 | 0.063 | 9.838 | **0.000** |
| Slope | -0.312 | 0.089 | -3.511 | **0.001** |
| **H.MFD** |  |  |  |  |
| intercept | 0.680 | 0.094 | 7.212 | **0.000** |
| Slope | -0.564 | 0.101 | -5.563 | **0.000** |

## Discussion

Our results demonstrate that neglect imperfect detection can produce unrealistic estimates of diversity, which can be unbalanced between treatment levels or environmental gradients. Considering that several community studies are pattern-based, ignoring the effect of imperfect detection can leads to spurious interpretations of the mechanisms driving community assembly (Joseph et al., 2009), mainly when inversion in the observed pattern occurs (critical bias regions, Fig. 1c). For the assemblage of fruit-feeding butterflies studied, we found a noise for site scale (the majority of points are in the noise region, Fig. 3b), typically produced by the inclusion of redundant species at understory for phylogenetic measures and redundant species at the canopy for functional measures. This occurs because the capacity to detect distinct lineages or functional traits in both strata was higher than the ability to detect new species (Jarzyna & Jetz, 2016), leading to an increase in phylogenetic or functional clustering in relation to the observed data. However, since between strata there is a difference in detection of individuals (reached by hidden diversity), the relationship between diversity and environment is biased.

Canopy and understory have distinct features including microclimatic conditions, forest structure, and resource availability (Grimbacher & Stork, 2007; Sobek, Tscharntke, Scherber, Schiele, & Steffan-Dewenter, 2009). Such differences are commonly associated with the vertical stratification of animal taxa, especially for insects (Ashton et al., 2016; Basset et al., 2015). For fruit-feeding butterflies, is recognized that some families or tribes are more related with canopy or understory (DeVries et al., 2012; Hill, Hamer, Tangah, & Dawood, 2001), including the detection probability of species can differ among strata (Ribeiro et al., 2016). Despite the lack of studies that access the phylogenetic and functional diversity for this group, and hence the relationship with forests strata, for the Neotropical region there is no clear pattern as to which is the most diverse stratum from a taxonomic perspective (understory – Araujo et al., 2020; Barlow, Overal, Araujo, Gardner, & Peres, 2007; Lourenço et al., 2019; Whitworth et al., 2016; canopy – Devries, 1988; DeVries et al., 2012; Ribeiro & Freitas, 2012; Santos et al., 2017). In our study, we show that there was a large underestimation in species richness, providing evidence that there is a bias for observed taxonomic diversity in canopies sites. This was the only case where there was an inversion in the observed pattern: understory was richest than canopy employing the observed data, but the canopy has higher richness than understory when we used the estimated data (Appendix C – Fig. C1, Table C1). For phylogenetic measures, despite the difference in HD values between stratum, the observed pattern was maintained and only the magnitude of the effect was adjusted. However, for functional measures based on distances, the inclusion of undetected individuals revealed a significant difference (understory was more diverse than canopy), unobserved when only observed data was used (Appendix C – Table C1).

As expected, the inclusion of undetected species had a larger effect on taxonomic diversity measures than on phylogenetic or functional ones. While for taxonomic diversity, each undetected species leads to an increment of the estimated diversity value, for phylogenetic and functional measures undetected species may be redundant, i.e., contain evolutionary or functional information, respectively, that was already covered in the observed data. Furthermore, we observed that the understory had a large number of species belonging to the same lineage that were undetected in the field. Generally, fruit-feeding butterflies that inhabit the understory belong to Satyrinae (particularly to the tribes Morphini and Brassolini). These species tend to be more abundant during the summer months (December to February in Southern Hemisphere) (Iserhard et al., 2017), and hence more individuals are available to be detected. But at the beginning or end of this season, a smaller number of individuals are active, hindering its detection. Such features could explain the clustered pattern observed in the understory when we include imperfect detection to perform phylogenetic measures. Similarly, a clustered pattern was revealed for functional measures for canopy. Species that occupy this stratum generally exhibit traits related to flight performance (Chai & Srygley, 1990; Graça et al., 2017), given high mobility to looking for resources and favorable conditions (Shahabuddin & Ponte, 2005). A simple explanation for the inclusion of redundant traits at canopy could be that individuals were absent because there were visiting a portion of this home range that was not covered by the survey (Joseph et al., 2009).

Biodiversity measures are important tools to guide species conservation decisions, as well as to infer about the ecological and evolutionary process that structure assemblages. Since accounting for imperfect detection improves the accuracy of estimates of diversity patterns, in some circumstances, it is strongly recommended (Fig 1), because it may lower the risk of erroneously inferring biological processes that are implied by sampling uncertainty (Joseph et al., 2009). Several models have been proposed in recent years to incorporate imperfect detection in order to improve the efficiency of estimating parameters in community studies (Abrams, Sollmann, Mitchell, Struebig, & Wilting, 2021; Broms et al., 2015; Frishkoff et al., 2017; Jarzyna & Jetz, 2016; Tingley, Nadeau, & Sandor, 2020; Zipkin et al., 2010). Further, these models allow us to propagate the uncertainty in species-specific detectability to biodiversity measures, as we demonstrated here. We expect that the framework developed in this study help researchers to better understand and describe diversity patterns and the mechanisms that assembly ecological communities.

## Acknowledgments

AR thanks Vivian Zulian and Gonçalo Ferraz for discussions about hierarchical models. AR thanks colleagues that helped with the butterfly surveys. The authors thank the staff of Floresta Nacional de São Francisco de Paula for providing logistic assistance during the sampling periods. AR research activities have been supported by Coordenação de Aperfeiçoamento de Pessoal de Nível Superior (CAPES) (graduate fellowship). LSD research activities have been supported by CNPq Productivity Fellowship (grant 307527/2018-2). AR, GN, CAI and LSD are members of the National Institutes for Science and Technology (INCT) in Ecology, Evolution and Biodiversity Conservation, supported by MCTIC/CNPq (proc. 465610/2014-5) and FAPEG (proc. 201810267000023).

