## Appendices A, B, and C for "The Hidden Side of Diversity: Effects of Imperfect Detection on Multiple Dimensions of Biodiversity": Appendix_A.docx

Code for the hierarchical Bayesian N-Mixture model (Static_Nmixture_P_RE.txt) and visual evaluation of parameters.

**N-mixture model:** This model is a modification of the proposed model for The Swiss Breeding Bird Survey (MHB 2014) in the Chapter 11 of the Applied Hierarchical Modelling in Ecology (Kéry & Royle, 2016). Furthermore, we assumes that the species-level effects can varies between strata, according whit Zipkin, Andrew Royle, Dawson, & Bates (2010). The code for the model is available on <https://github.com/richterbine/TheHiddenSideofDiversity/tree/main/R/Bayesian_models>

model {

\# Community priors (with hyperparameters) for species-specific parameters

for(k in 1:nspec){

beta.can[k] ~ dnorm(mu.beta.can, tau.beta.can) # Species-specific random intercept for abundance - canopy

beta.und[k] ~ dnorm(mu.beta.und, tau.beta.und) # Species-specific random intercept for abundance - understory

alpha.can[k] ~ dnorm(mu.alpha.can, tau.alpha.can) # Species-specific random intercept for detectability - canopy

alpha.und[k] ~ dnorm(mu.alpha.und, tau.alpha.und) # Species-specific random intercept for detectability - understory

beta1[k] ~ dnorm(mu.beta1, tau.beta1) # Species-specific random slope for temperature in biologic process

alpha1[k] ~ dnorm(mu.alpha1, tau.alpha1) # Species-specific random slope for sampling process in sampling process

alpha2[k] ~ dnorm(mu.alpha2, tau.alpha2) # Species-specific random slope for temperature in sampling process

for(n in 1:5) {

month[n, k] ~ dnorm(0, tau.month)

}

for(n in 1:6){

area[n, k] ~ dnorm(0, tau.area)

}

}

\# Hyperpriors for community hyperparameters

\# abundance model - intercept

mu.beta.can ~ dnorm(0, 0.001)

tau.beta.can <- pow(sd.beta.can, -2)

sd.beta.can ~ dunif(0, 10)

mu.beta.und ~ dnorm(0, 0.001)

tau.beta.und <- pow(sd.beta.und, -2)

sd.beta.und ~ dunif(0, 10)

\# abundance model - slope for temperature by sites

mu.beta1 ~ dnorm(0, 0.001)

tau.beta1 <- pow(sd.beta1, -2)

sd.beta1 ~ dunif(0, 10)

\# detection model - intercept

mu.alpha.can.pre ~ dunif (0, 1) # Detection can have any value between 0 and 1 with equal probability

mu.alpha.can <- logit(mu.alpha.can.pre) # Inverse logit – values from -inf to inf as in norm

tau.alpha.can <- pow(sd.alpha.can, -2)

sd.alpha.can ~ dunif(0, 10)

mu.alpha.und.pre ~ dunif(0, 1)

mu.alpha.und <- logit(mu.alpha.und.pre)

tau.alpha.und <- pow(sd.alpha.und, -2)

sd.alpha.und ~ dunif(0, 10) ## sd

\# detection model - slope for julian dates

mu.alpha1 ~ dnorm(0, 0.001)

tau.alpha1 <- pow(sd.alpha1, -2)

sd.alpha1 ~ dunif(0, 10)

\# detection model - slope for temperatures per day

mu.alpha2 ~ dnorm(0, 0.001)

tau.alpha2 <- pow(sd.alpha2, -2)

sd.alpha2 ~ dunif(0, 10)

tau.month <- pow(sd.month, -2)

sd.month ~ dunif(0, 10)

tau.area <- pow(sd.area, -2)

sd.area ~ dunif(0, 10)

\# Ecological model for true abundance (process model)

for(k in 1:nspec) {

for (i in 1:nsite) {

N[i,k] ~ dpois(lambda[i,k])	# latent abundance of each species in each site

log(lambda[i,k]) <- beta.can[k] * (1 - Strata[i]) + beta.und[k] * Strata[i] + beta1[k] * Temp[i] + month[Month[i], k] + area[Area[i], k]

}

}

\# Observation model for replicated counts

for(k in 1:nspec) {

for (i in 1:nsite) {

for (j in 1:nrep) {

yc[i,j,k] ~ dbin(p[i,j,k], N[i,k])

logit(p[i,j,k]) <- alpha.can[k] * (1 - Strata[i]) + alpha.und[k] * Strata[i] + alpha1[k] * Date[i, j] + alpha2[k] * Temp_det[i, j]

}

}

}

\# Other derived quantities

for(k in 1:nspec) {

mlambda.can[k] <- exp(beta.can[k]) # Expected abundance on natural scale for canopy

mlambda.und[k] <- exp(beta.und[k]) # Expected abundance on natural scale for understory

logit(mp.can[k]) <- alpha.can[k]	# Mean detection on natural scale for canopy

logit(mp.und[k]) <- alpha.und[k] # Mean detection on natural scale for understory

}

}

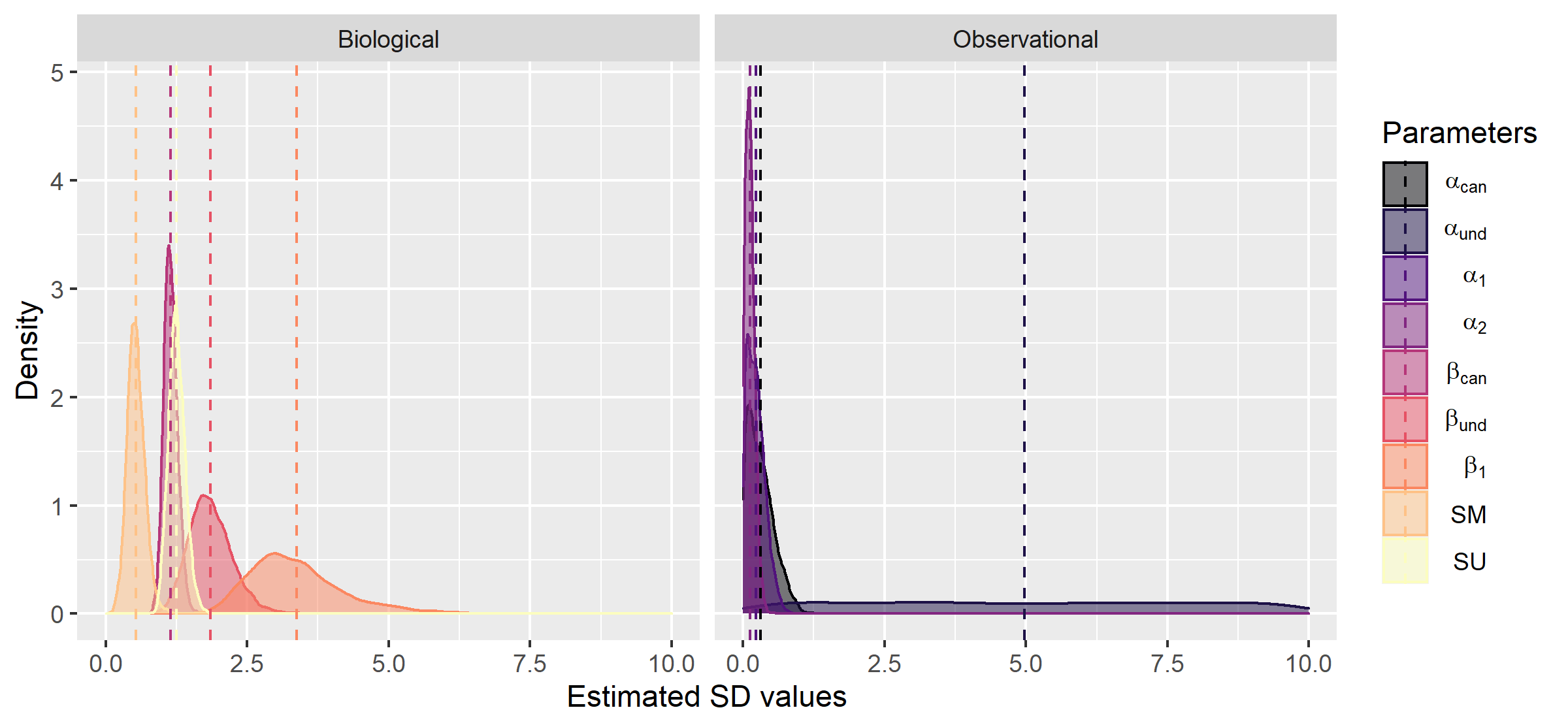

**Figure A 1**. Community distribution of the standard deviation for mean parameters, estimated by N-mixture models for biological and observational processes, using observed data for fruit-feeding butterflies communities sampled at FLONA-SFP, Southern Brazil.

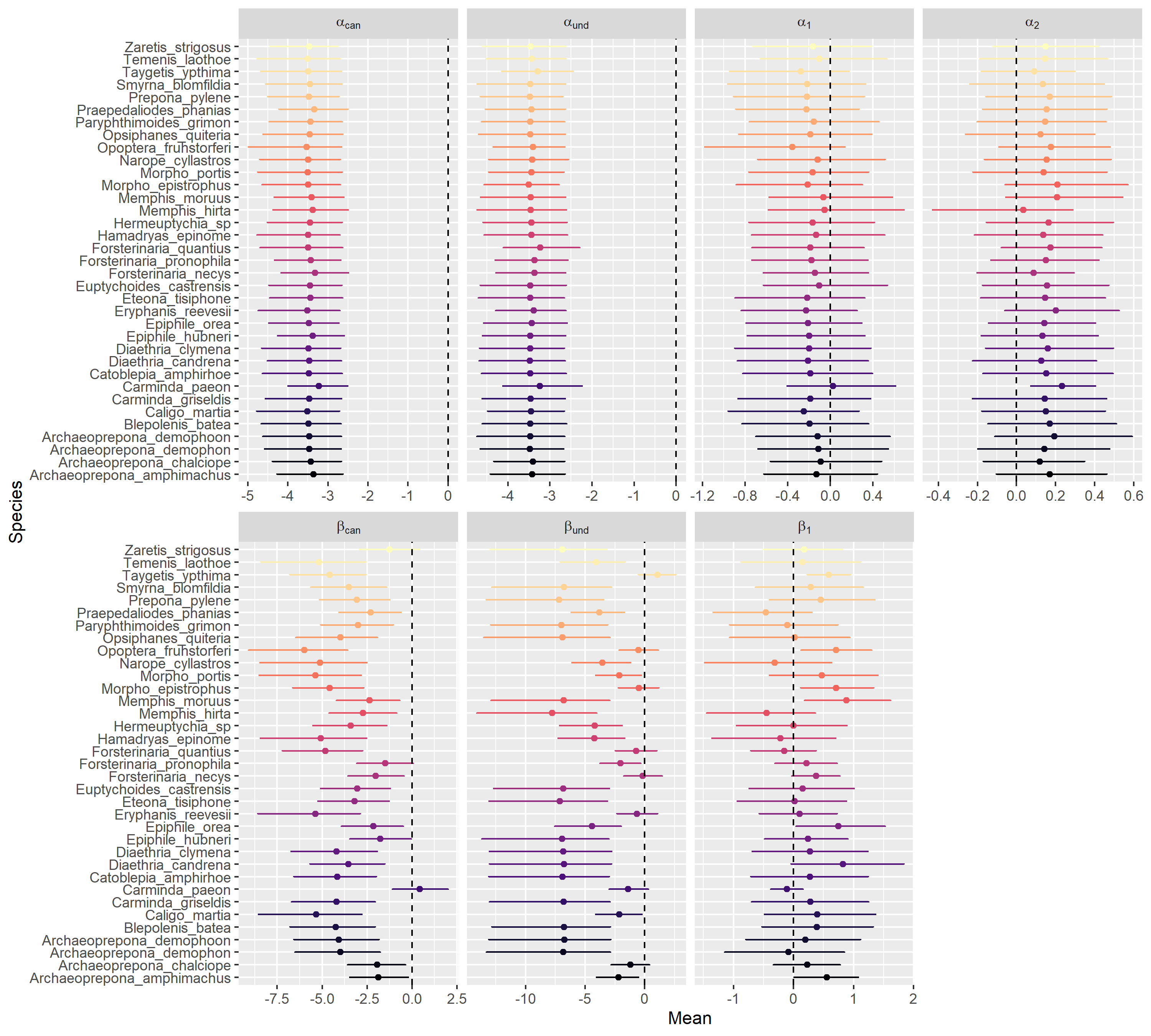

**Figure A 2**. Specie-specific mean parameters were estimated by the N-mixture model for detection probability (upper panel) and abundance (lower panel). Points represent the estimated parameter for each species and horizontal bars indicate the 95% of credible interval (CRI). If the CRI overlaps the zero (vertical dashed line), we conclude that there was no effect of the predictor in species-specific responses. For detection probability, sampling day (α_1_) does not affect the detection of any species, while the temperature at trap base (α_2_) only affected positively the species *Carminda paeon* (Godart, 1824). For expected abundance, temperature (β_1_) has a positive effect for *Taygetis ypthima* Hübner, [1821], *Opoptera fruhstorferi* (Röber, 1896), *Morpho epistrophus* (Fabricius, 1796), *Memphis moruus* (Fabricius, 1775), *Epiphile orea* (Hübner, [1823]).

**Table A 1**. The mean and the 95% Bayesian credible interval (CRI) for community-level summaries of the hyper-parameters for expected abundance (β’s) and detection probability (α’s) estimated by the N-mixture model for fruit-feeding butterflies sampled in FLONA-SFP, southern Brazil. Asterisk represents hyper-parameters that overlap zero.

|  | Mean | Low CRI | Upper CRI |
| --- | --- | --- | --- |
| µ β_can_ | -3.4991 | -4.4792 | -2.5917 |
| SD β_can_ | 1.8470 | 1.1903 | 2.6484 |
| µ β_und_ | -4.4248 | -6.3377 | -2.9100 |
| SD β_und_ | 3.3726 | 2.1065 | 5.3857 |
| µ β_1_ | 0.2237 | -0.0740 | 0.4975* |
| SD β_1_ | 0.5334 | 0.2564 | 0.8592 |
| µα_can_ | -3.4379 | -4.0621 | -2.9272 |
| SD α_can_ | 0.3105 | 0.0097 | 0.8413 |
| µ α_und_ | -0.0359 | -3.9749 | 3.6357* |
| SD α_und_ | 4.9700 | 0.2257 | 9.7353 |
| µα_1_ | -0.1693 | -0.4526 | 0.1057* |
| SD α_1_ | 0.2276 | 0.0118 | 0.5851 |
| µα_2_ | 0.1520 | 0.0094 | 0.2878 |
| SD α_2_ | 0.1233 | 0.0048 | 0.3062 |
| SD SM | 1.2553 | 0.9940 | 1.5692 |
| SD SU | 1.1465 | 0.9365 | 1.3983 |
