## Appendices A, B, and C for "The Hidden Side of Diversity: Effects of Imperfect Detection on Multiple Dimensions of Biodiversity": Appendix_B.docx

Visual evaluation of the performance of the N-mixture model built. We used 12 simulated communities, which we vary parameters related to biological process (β_1_ and β_2_) and observational process (α_1_ and α_2_). We consider that model is valid if the values of the real parameters are within the posterior distribution retuned by the N-mixture model. The function for simulate metacommunities is a modified version of the *simComm* function developed by Kéry & Royle (2016) and available in the AHMBook package (v. 0.2.2). The code to simulate metacommunities are available on <https://github.com/richterbine/TheHiddenSideofDiversity/tree/main/R/Model_validation>

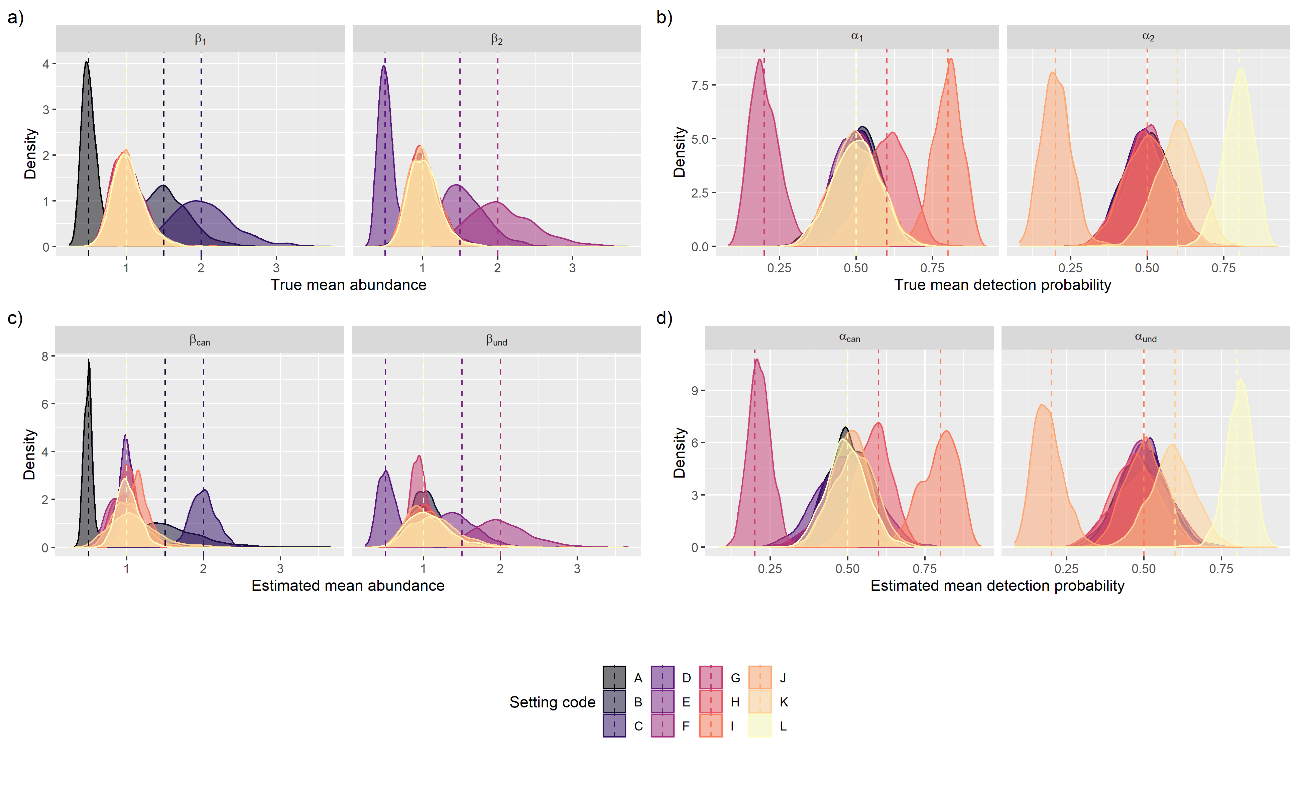

**Figure B 1**. Community distribution of mean abundance and detection probability for simulated communities. a) and b) are the distribution using the true values for intercept parameters (µ_β1_, sd_β1,_ µ_β2_, sd_β2,_ µ_α1_, sd_α1,_ µ_α2_, sd_α2_). c) and d) are the distribution using the mean values estimated by the Bayesian model (µ_βcan_, sd_βcan_, µ_αcan_, sd_αcan_, µ_βund_, sd_βund_, µ_αund_, sd_αund_). The color indicates the 12 scenarios simulated, when we vary the values for µ_β1,_ µ_β2,_ µ_α1,_ µ_α2_ for three levels: µ_β_ = 0.5, 1.5, 2.0, the default of function is 1.0; µ_α_ = 0.2, 0.6, 0.8, the default of function is 0.5. We allow only one parameter to vary at a time, while others remained fixed to simulate the community. For A, B and C the µ_β1_ varies, for D, E and F the µ_β2_ varies, for G, H and I the µ_α1_ varies and for J, K and L the µ_α2_ varies.

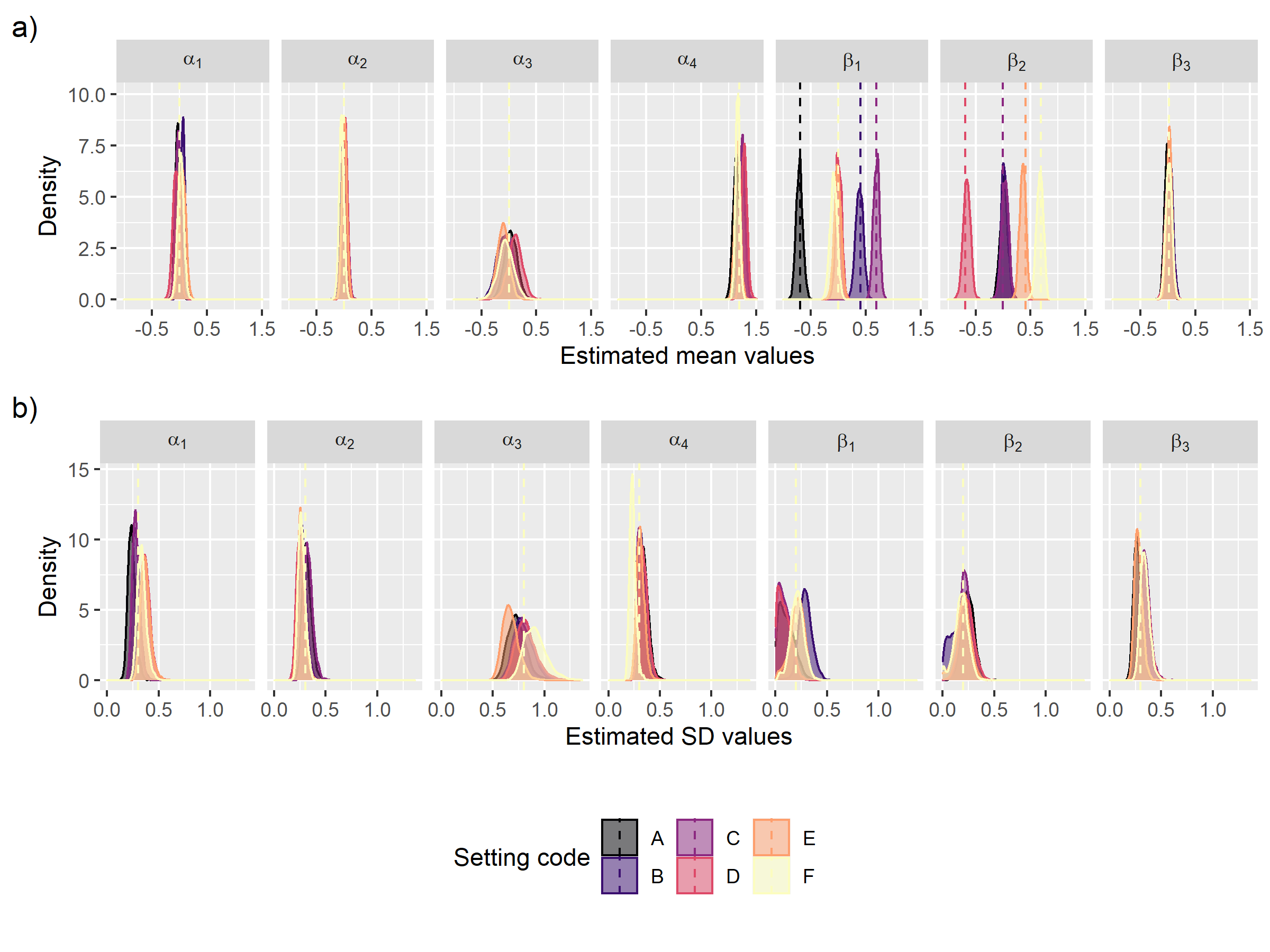

**Figure B 2**. Posterior distribution of estimated hyper-parameters (µ and sd) for simulated communities. a) and b) are the estimated mean values for all parameters considering the models simulated varying the mean abundance (β values, setting A to F). The dashed line represents the true values used to simulate the community for each set.

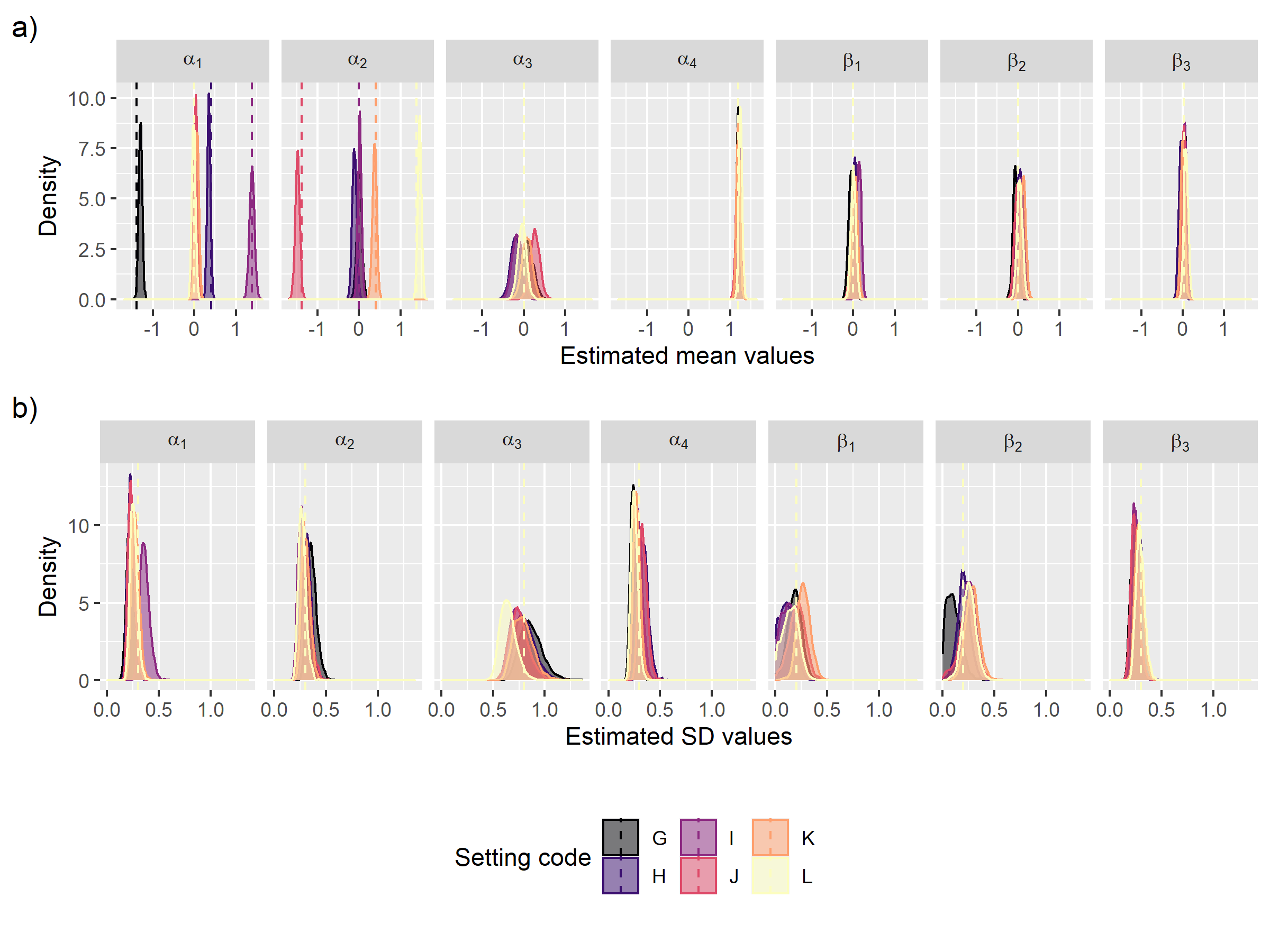

**Figure B 3**. Posterior distribution of estimated hyper-parameters (µ and sd) for simulated communities. a) and b) are the estimated values considering the communities simulated varying the mean detection probability (α values, setting G to L). The dashed line represents the true values used to simulate the community for each set.
