## Appendices A, B, and C for "The Hidden Side of Diversity: Effects of Imperfect Detection on Multiple Dimensions of Biodiversity": Appendix_C.docx

**Hidden diversity framework**: Function to evaluate the extent to which imperfect detection may affect patterns of taxonomic, functional, and phylogenetic diversities in ecological communities This function allows the user to estimate, in the form of a Standardized Effect Size (SES), how much of the diversity was hidden when we do not account with the imperfect detection. Further, the user can calculate the hidden diversity only for taxonomic diversity (TD and Abundance), for functional (SES.FD and SES.MFD), and taxonomic diversity, for phylogenetic (SES.PD and SES.MPD) and taxonomic diversity, or for all measures.

**Arguments**

comm = Community data, with sites in the rows and species in the columns.

N = an array when each “slice” is a matrix of the true-abundance (sites in rows and species in columns) for one posterior sampling of the N-mixture model. This object represents the detection-corrected abundance.

phy = a phylogenetic tree, with branch length.

trait = a trait community matrix, with species in rows and traits in columns.

binary = logical. Default is FALSE. Only necessary when a trait matrix is present. If binary = TRUE, the function is taken into account binary traits for construct a functional dendrogram.

abundance.weighted = logical. Default is FALSE. In this case, for SES.MPD and SES.MFD only the occurrence/incidence/richness of species is accounted by calculating the diversity values. If abundance.weighted = TRUE, the SES.MPD and SES.MFD will be calculated for an abundance-based community matrix.

Null.model = a string with the null models allowed by ses.pd and ses.mpd function from *picante* package.

Runs = number of permutations used to calculate the null models.

**Value**

The function returns a list with two, three, or four data frames (dependent on imputed diversity):

TD.df, N.df = always returned. Each object is an object with four columns: the observed richness/abundance, the mean richness/abundance for estimated data (N), the standard deviation of the estimated richness/abundance, and the hidden diversity.

PD.df, FD.df, MPD.df and MFD.df = only returned if the user informed a phylogenetic tree and/or a functional traits matrix. Each object has seven columns: the observed diversity value, the SES value for observed diversity, the mean estimated diversity value, the SES values for mean estimated diversity, the standard deviation for estimated diversity, the standard deviation for estimated SES.

hidden.diversity <- function(comm, N, phy = NULL, trait = NULL, binary = FALSE, abundance.weighted = FALSE, null.model = "taxa.labels", runs = 999) {

n.site <- dim(N)[1]	# n.site: the number of sampling sites

n.samp <- dim(N)[3]	# n.samp: the number of posterior sampling

\# transforming N in occurrence data (y)

y <- N

for (i in 1: dim(y)[3]) {

b = which(y[, , i] > 0)

y[, , i][b] = 1

y[, , i][-b] = 0

}

\# calculating the observed and estimated richness (TD)

TD.df <- data.frame(TD.obs = apply(vegan::decostand(x = comm, method = "pa"), 1, sum), TD.mean = apply(apply(y, c(1,3), sum), 1, mean), TD.sd = apply(apply(y, c(1,3), sum), 1, sd))

\# calculating the observed and estimated abundance

N.df <- data.frame(N.obs = apply(comm, 1, sum), N.mean = apply(apply(N, c(1,3), sum), 1, mean), N.sd = apply(apply(N, c(1,3), sum), 1, sd))

\# if only the phylogenetic tree is provided

if (is.null(trait) == TRUE & is.null(phy) == FALSE){

pd.ses <- array(NA, dim = c(n.site, 2, n.samp))

mpd.ses <- array(NA, dim = c(n.site, 2, n.samp))

dist.phylo <- cophenetic(x = phy)

\# calculating the phylogenetic diversity (SES.PD)

\# observed data

pd.obs <- picante::ses.pd(samp = comm, tree = phy, null.model = null.model, runs = runs, include.root = F)

\# estimated data

for (i in 1:n.samp){

temp <- picante::ses.pd(samp = N[, , i], tree = phy, null.model = null.model, runs = runs, include.root = F)

pd.ses[,1,i] <- cbind(temp[,2])

pd.ses[,2,i] <- cbind(temp[,6])

}

PD.df <- data.frame(PD.obs = pd.obs[, "pd.obs"], SES.PD.obs = pd.obs[, "pd.obs.z"], PD.est = apply(pd.ses[,1,], 1, mean, na.rm = T),

SES.PD.est = apply(pd.ses[,2,], 1, mean, na.rm = T), PD.sd = apply(pd.ses[,1,], 1, sd, na.rm = T),

SES.PD.sd = apply(pd.ses[,2,], 1, sd, na.rm = T))

\## calculating the mean phylogenetic diversity (SES.MPD)

\# observed data

mpd.obs <- picante::ses.mpd(samp = comm, dis = dist.phylo, abundance.weighted = abundance.weighted, null.model = null.model, runs = runs)

\# estimated data

for (i in 1:n.samp){

temp <- picante::ses.mpd(samp = N[, , i], dis = dist.phylo, abundance.weighted = abundance.weighted,

null.model = null.model, runs=runs)

mpd.ses[,1,i] <- cbind(temp[,2])

mpd.ses[,2,i] <- cbind(temp[,6])

}

MPD.df <- data.frame(MPD.obs = pd.obs[, "pd.obs"], SES.MPD.obs = pd.obs[, "pd.obs.z"],

MPD.est = apply(mpd.ses[,1,], 1, mean, na.rm = T), SES.MPD.est = apply(mpd.ses[,2,], 1, mean, na.rm = T),

MPD.sd = apply(mpd.ses[,1,], 1, sd, na.rm = T), SES.MPD.sd = apply(mpd.ses[,2,], 1, sd, na.rm = T))

\# calculating the hidden diversity

TD.df$hidden.TD <- (TD.df$TD.obs - TD.df$TD.mean) / TD.df$TD.sd

N.df$hidden.N <- (N.df$N.obs - N.df$N.mean) / N.df$N.sd

PD.df$hidden.PD <- (PD.df$SES.PD.obs - PD.df$SES.PD.est) / PD.df$SES.PD.sd

MPD.df$hidden.MPD <- (MPD.df$SES.MPD.obs - MPD.df$SES.MPD.est) / MPD.df$SES.MPD.sd

return(list(TD.df, N.df, PD.df, MPD.df))

}

\# if only a traits matrix is provided

if (is.null(trait) == FALSE & is.null(phy) == TRUE){

if(binary == TRUE){

bin <- vector()

for(i in 1:ncol(trait)){

bin[i] <- is.integer(trait[, i]) | is.factor(trait[, i])

}

con.t <- which(bin == F)

bin.t <- which(bin == T)

t.dist <- ade4::dist.ktab(ade4::ktab.list.df(list(log(trait[, con.t]), ade4::prep.binary(trait[, bin.t], col.blocks = ncol(trait[, bin.t])))),

type = c("Q", "B")) # create a dist matrix, considering mixed-variables

} else {

t.dist <- ade4::dist.ktab(ade4::ktab.list.df(list(log(trait))), type = "Q")

}

tree.func <- hclust(d = t.dist, method = "average") # clustering using UPGMA

tree.func <- ape::as.phylo(tree.func)

fd.ses <- array(NA, dim = c(n.site, 2, n.samp))

mfd.ses <- array(NA, dim = c(n.site, 2, n.samp))

dist.func <- cophenetic(tree.func)

\# calculating the functional diversity (SES.FD)

\# observed data

fd.obs <- picante::ses.pd(samp = comm, tree = tree.func, null.model = null.model, runs = runs, include.root = F)

\# estimated data

for (i in 1:n.samp){

temp <- picante::ses.pd(samp = N[,,i], tree = tree.func, null.model = null.model, runs = runs, include.root = F)

fd.ses[,1,i] <- cbind(temp[,2])

fd.ses[,2,i] <- cbind(temp[,6])

}

FD.df <- data.frame(FD.obs = fd.obs[, "pd.obs"], SES.FD.obs = fd.obs[, "pd.obs.z"], FD.est = apply(fd.ses[,1,], 1, mean, na.rm = T),

SES.FD.est = apply(fd.ses[,2,], 1, mean, na.rm = T), FD.sd = apply(fd.ses[,1,], 1, sd, na.rm = T),

SES.FD.sd = apply(fd.ses[,2,], 1, sd, na.rm = T))

\## calculating the mean functional diversity (SES.MFD)

\# observed data

mfd.obs <- picante::ses.mpd(samp = comm, dis = dist.func, null.model = null.model, abundance.weighted = abundance.weighted,

runs = runs)

\# estimated data

for (i in 1:n.samp){

temp <- picante::ses.mpd(samp = N[, , i], dis = dist.func, null.model = null.model, abundance.weighted = abundance.weighted, runs = runs)

mfd.ses[,1,i] <- cbind(temp[,2])

mfd.ses[,2,i] <- cbind(temp[,6])

}

MFD.df <- data.frame(MFD.obs = mfd.obs[, "mpd.obs"], SES.MFD.obs = mfd.obs[, "mpd.obs.z"],

MFD.est = apply(mfd.ses[,1,], 1, mean, na.rm = T), SES.MFD.est = apply(mfd.ses[,2,], 1, mean, na.rm = T),

MFD.sd = apply(mfd.ses[,1,], 1, sd, na.rm = T), SES.MFD.sd = apply(mfd.ses[,2,], 1, sd, na.rm = T))

\# calculating the hidden diversity

TD.df$hidden.TD <- (TD.df$TD.obs - TD.df$TD.mean) / TD.df$TD.sd

N.df$hidden.N <- (N.df$N.obs - N.df$N.mean) / N.df$N.sd

FD.df$hidden.FD <- (FD.df$SES.FD.obs - FD.df$SES.FD.est) / FD.df$SES.FD.sd

MFD.df$hidden.MFD <- (MFD.df$SES.MFD.obs - MFD.df$SES.MFD.est) / MFD.df$SES.MFD.sd

return(list(TD.df, N.df, FD.df, MFD.df))

}

\# if both phylogenetic tree and traits matrix are provided

if (is.null(trait) == FALSE & is.null(phy) == FALSE){

pd.ses <- array(NA, dim = c(n.site, 2, n.samp))

mpd.ses <- array(NA, dim = c(n.site, 2, n.samp))

dist.phylo <- cophenetic(x = phy)

\# calculating the phylogenetic diversity (SES.PD)

\# observed data

pd.obs <- picante::ses.pd(samp = comm, tree = phy, null.model = null.model, runs = runs, include.root = F)

\# estimated data

for (i in 1:n.samp){

temp <- picante::ses.pd(samp = N[,,i], tree = phy, null.model = null.model, runs = runs, include.root = F)

pd.ses[,1,i] <- cbind(temp[,2])

pd.ses[,2,i] <- cbind(temp[,6])

}

PD.df <- data.frame(PD.obs = pd.obs[, "pd.obs"], SES.PD.obs = pd.obs[, "pd.obs.z"], PD.est = apply(pd.ses[,1,], 1, mean, na.rm = T),

SES.PD.est = apply(pd.ses[,2,], 1, mean, na.rm = T), PD.sd = apply(pd.ses[,1,], 1, sd, na.rm = T),

SES.PD.sd = apply(pd.ses[,2,], 1, sd, na.rm = T))

\## calculating the mean phylogenetic diversity (SES.MPD)

\# observed data

mpd.obs <- picante::ses.mpd(samp = comm, dis = dist.phylo, abundance.weighted = abundance.weighted, null.model = null.model, runs = runs)

\# estimated data

for (i in 1:n.samp){

temp <- picante::ses.mpd(samp = N[, , i], dis = dist.phylo, abundance.weighted = abundance.weighted, null.model = null.model, runs=runs)

mpd.ses[,1,i] <- cbind(temp[,2])

mpd.ses[,2,i] <- cbind(temp[,6])

}

MPD.df <- data.frame(MPD.obs = pd.obs[, "pd.obs"], SES.MPD.obs = pd.obs[, "pd.obs.z"],

MPD.est = apply(mpd.ses[,1,], 1, mean, na.rm = T), SES.MPD.est = apply(mpd.ses[,2,], 1, mean, na.rm = T),

MPD.sd = apply(mpd.ses[,1,], 1, sd, na.rm = T), SES.MPD.sd = apply(mpd.ses[,2,], 1, sd, na.rm = T))

\# Calculating the functional diversity

if(binary == TRUE){

bin <- vector()

for(i in 1:ncol(trait)){

bin[i] <- is.integer(trait[, i]) | is.factor(trait[, i])

}

con.t <- which(bin == F)

bin.t <- which(bin == T)

t.dist <- ade4::dist.ktab(ade4::ktab.list.df(list(log(trait[, con.t]), ade4::prep.binary(trait[, bin.t], col.blocks = ncol(trait[, bin.t])))),

type = c("Q", "B")) # create a dist matrix, considering mixed-variables

} else {

t.dist <- ade4::dist.ktab(ade4::ktab.list.df(list(log(trait))), type = "Q")

}

tree.func <- hclust(d = t.dist, method = "average") # clustering using UPGMA

tree.func <- ape::as.phylo(tree.func)

fd.ses <- array(NA, dim = c(n.site, 2, n.samp))

mfd.ses <- array(NA, dim = c(n.site, 2, n.samp))

dist.func <- cophenetic(tree.func)

\# calculating the functional diversity (SES.FD)

\# observed data

fd.obs <- picante::ses.pd(samp = comm, tree = tree.func, null.model = null.model, runs = runs, include.root = F)

\# estimated data

for (i in 1:n.samp){

temp <- picante::ses.pd(samp = N[, , i], tree = tree.func, null.model = null.model, runs = runs, include.root = F)

fd.ses[,1,i] <- cbind(temp[,2])

fd.ses[,2,i] <- cbind(temp[,6])

}

FD.df <- data.frame(FD.obs = fd.obs[, "pd.obs"], SES.FD.obs = fd.obs[, "pd.obs.z"], FD.est = apply(fd.ses[,1,], 1, mean, na.rm = T),

SES.FD.est = apply(fd.ses[,2,], 1, mean, na.rm = T), FD.sd = apply(fd.ses[,1,], 1, sd, na.rm = T),

SES.FD.sd = apply(fd.ses[,2,], 1, sd, na.rm = T))

\## calculating the mean functional diversity (SES.MFD)

\# observed data

mfd.obs <- picante::ses.mpd(samp = comm, dis = dist.func, null.model = null.model, abundance.weighted = abundance.weighted,

runs = runs)

\# estimated data

for (i in 1:n.samp){

temp <- picante::ses.mpd(samp = N[, , i], dis = dist.func, null.model = null.model, abundance.weighted = abundance.weighted, runs = runs)

mfd.ses[,1,i] <- cbind(temp[,2])

mfd.ses[,2,i] <- cbind(temp[,6])

}

MFD.df <- data.frame(MFD.obs = mfd.obs[, "mpd.obs"], SES.MFD.obs = mfd.obs[, "mpd.obs.z"],

MFD.est = apply(mfd.ses[,1,], 1, mean, na.rm = T), SES.MFD.est = apply(mfd.ses[,2,], 1, mean, na.rm = T),

MFD.sd = apply(mfd.ses[,1,], 1, sd, na.rm = T), SES.MFD.sd = apply(mfd.ses[,2,], 1, sd, na.rm = T))

\# calculating the hidden diversity

TD.df$hidden.TD <- (TD.df$TD.obs - TD.df$TD.mean) / TD.df$TD.sd

N.df$hidden.N <- (N.df$N.obs - N.df$N.mean) / N.df$N.sd

PD.df$hidden.PD <- (PD.df$SES.PD.obs - PD.df$SES.PD.est) / PD.df$SES.PD.sd

MPD.df$hidden.MPD <- (MPD.df$SES.MPD.obs - MPD.df$SES.MPD.est) / MPD.df$SES.MPD.sd

FD.df$hidden.FD <- (FD.df$SES.FD.obs - FD.df$SES.FD.est) / FD.df$SES.FD.sd

MFD.df$hidden.MFD <- (MFD.df$SES.MFD.obs - MFD.df$SES.MFD.est) / MFD.df$SES.MFD.sd

return(list(TD.df, N.df, PD.df, MPD.df, FD.df, MFD.df))

}

\# if neither the phylogenetic tree nor the traits matrix is provided

else {

\## calculating the hidden diversity

TD.df$hidden.TD <- (TD.df$TD.obs - TD.df$TD.mean) / TD.df$TD.sd

N.df$hidden.N <- (N.df$N.obs - N.df$N.mean) / N.df$N.sd

return(list(TD.df, N.df))

}

}

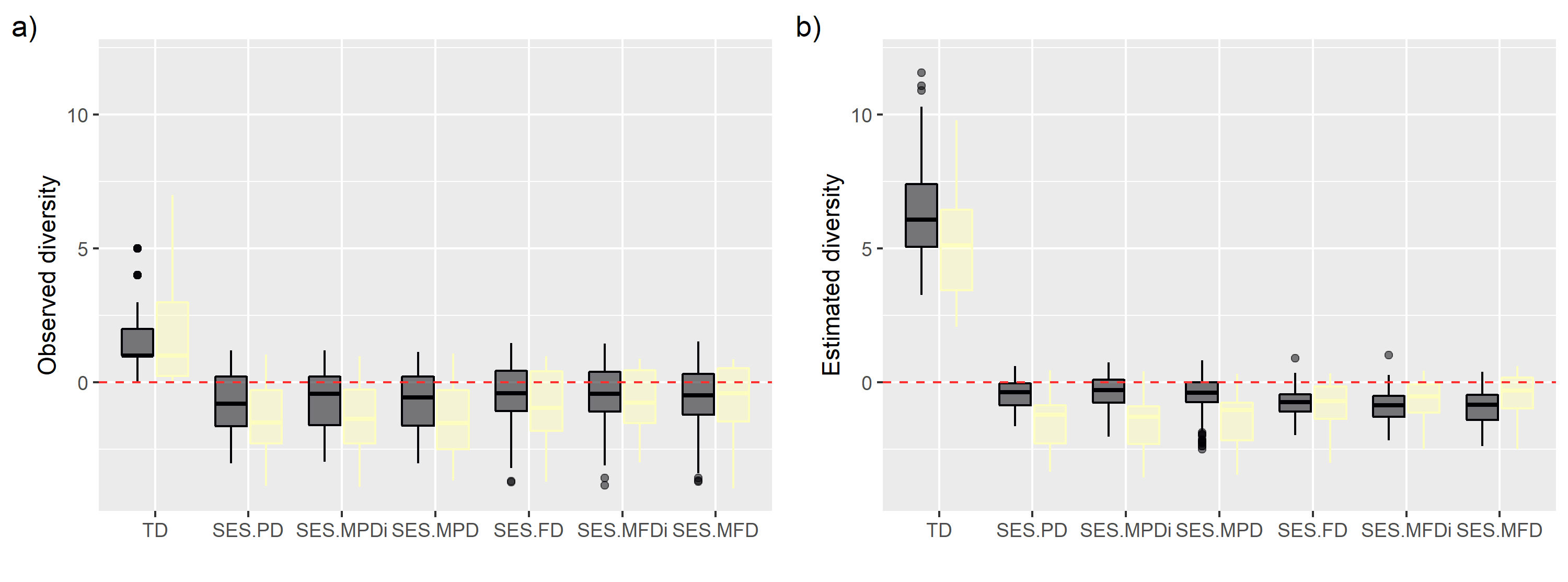

**Figure C 1**. Relationship among diversity measures and canopy (dark boxplots) and understory (yellow boxplots). a) Diversity pattern calculated using only observed data. b) Diversity pattern calculated using 100 matrices of estimated true abundance. TD – taxonomic diversity, SES – standardized effect size, PD/FD – phylogenetic/functional diversity, MPD/MFD – abundance-based mean pairwise phylogenetic/functional distance, MPDi/MFDi – incidence-based mean pairwise phylogenetic/functional distance.

**Table C 1**. Relationship between distinct facets of biodiversity and vertical stratification for fruit-feeding butterflies sampled at FLONA-SFP, southern Brazil. The first four columns show the relationship of diversity measures obtained by observed data with canopy and understory, and the last four columns show the relationship of diversity measures obtained by estimated data (corrected-by-detection). Bold values indicate a statistical significance at a threshold of 0.05. Asterisk indicates the unique case where there was an inversion of the most diverse stratum. TD – taxonomic diversity, SES – standardized effect size, PD/FD – phylogenetic/functional diversity, MPD/MFD – abundance-based mean pairwise phylogenetic/functional distance, MPDi/MFDi – incidence-based mean pairwise phylogenetic/functional distance.

|  | Observed data | | | |  | Estimated data | | | |
| --- | --- | --- | --- | --- | --- | --- | --- | --- | --- |
|  | Estimate | SE | t value | p |  | Estimate | SE | t value | p |
| TD |  |  |  |  |  |  |  |  |  |
| Canopy | 1.440 | 0.373 | 3.860 | **0.004** |  | 6.296 | 0.785 | 8.021 | **0.000** |
| Understory* | 0.280 | 0.140 | 2.003 | **0.046** |  | -1.169 | 0.088 | -13.35 | **0.000** |
| SES.PD |  |  |  |  |  |  |  |  |  |
| Canopy | -0.547 | 0.297 | -1.841 | 0.108 |  | -0.459 | 0.264 | -1.739 | 0.132 |
| Understory | -0.584 | 0.194 | -3.003 | **0.003** |  | -1.017 | 0.048 | -21.33 | **0.000** |
| SES.FD |  |  |  |  |  |  |  |  |  |
| Canopy | -0.589 | 0.315 | -1.873 | 0.094 |  | -0.772 | 0.213 | -3.619 | **0.009** |
| Understory | 0.009 | 0.200 | 0.047 | 0.963 |  | -0.038 | 0.049 | -0.764 | 0.446 |
| SES.MPDi |  |  |  |  |  |  |  |  |  |
| Canopy | -0.506 | 0.304 | -1.668 | 0.137 |  | -0.382 | 0.277 | -1.379 | 0.212 |
| Understory | -0.580 | 0.186 | -3.118 | **0.002** |  | -1.140 | 0.051 | -22.29 | **0.000** |
| SES.MPD |  |  |  |  |  |  |  |  |  |
| Canopy | -0.546 | 0.318 | -1.719 | 0.128 |  | -0.480 | 0.315 | -1.523 | 0.172 |
| Understory | -0.568 | 0.184 | -3.084 | **0.003** |  | -0.930 | 0.050 | -18.48 | **0.000** |
| SES.MFDi |  |  |  |  |  |  |  |  |  |
| Canopy | -0.626 | 0.306 | -2.044 | 0.072 |  | -0.914 | 0.218 | -4.199 | **0.004** |
| Understory | 0.087 | 0.201 | 0.433 | 0.666 |  | 0.276 | 0.045 | 6.188 | **0.000** |
| SES.MFD |  |  |  |  |  |  |  |  |  |
| Canopy | -0.680 | 0.341 | -1.996 | 0.076 |  | -0.922 | 0.263 | -3.505 | **0.008** |
| Understory | 0.175 | 0.202 | 0.866 | 0.388 |  | 0.463 | 0.044 | 10.61 | **0.000** |
